# TRIM46 is not required for axon specification or axon initial segment formation *in vivo*

**DOI:** 10.1101/2024.05.23.595556

**Authors:** Allison J. Melton, Victoria L. Palfini, Yuki Ogawa, Matthew N. Rasband

**Affiliations:** Department of Neuroscience, Baylor College of Medicine, Houston, TX, USA 77030

**Keywords:** TRIM46, axon initial segment, AnkG, microtubule fasciculation, node of Ranvier

## Abstract

Vertebrate nervous systems use the axon initial segment (AIS) to initiate action potentials and maintain neuronal polarity. The microtubule-associated protein tripartite motif containing 46 (TRIM46) was reported to regulate axon specification, AIS assembly, and neuronal polarity through the bundling of microtubules in the proximal axon. However, these claims are based on TRIM46 knockdown in cultured neurons. To investigate TRIM46 function *in vivo*, we examined TRIM46 knockout mice. Contrary to previous reports, we find that TRIM46 is dispensable for AIS formation and maintenance, and axon specification. TRIM46 knockout mice are viable, have normal behavior, and have normal brain structure. Thus, TRIM46 is not required for AIS formation, axon specification, or nervous system function. We also show TRIM46 enrichment in the first ∼100 μm of axon occurs independently of ankyrinG (AnkG), although AnkG is required to restrict TRIM46 only to the AIS. Our results suggest an unidentified protein may compensate for loss of TRIM46 *in vivo* and highlight the need for further investigation of the mechanisms by which the AIS and microtubules interact to shape neuronal structure and function.

**SIGNIFICANCE STATEMENT:** A healthy nervous system requires the polarization of neurons into structurally and functionally distinct compartments, which depends on both the axon initial segment (AIS) and the microtubule cytoskeleton. In contrast to previous reports, we show that the microtubule-associated protein TRIM46 is not required for axon specification or AIS formation in mice. Our results emphasize the need for further investigation of the mechanisms by which the AIS and microtubules interact to shape neuronal structure and function.

## INTRODUCTION

Vertebrate nervous systems depend on the axon initial segment (AIS) for both action potential firing and the maintenance of neuronal polarity (Leterrier, 2018). The AIS was first identified over 50 years ago on the basis of several ultrastructural features visible with electron microscopy (Palay et al., 1968; Peters et al., 1968). Of these, the best understood is an electron-dense material lining the plasma membrane. This material consists of tightly packed voltage-gated ion channels and cell adhesion molecules that are recruited and anchored to the submembranous actin-spectrin cytoskeleton by the master organizer of the AIS: the scaffolding protein ankyrinG (AnkG). Loss of AnkG abolishes AIS formation, impairs action potential firing, disrupts the structural and molecular characteristics of the axon, and is fatal at birth (Zhou et al., 1998; Hedstrom et al., 2008; Jenkins et al., 2013, 2015; Ho et al., 2014). Therefore, investigation of the electron-dense material lining the AIS membrane has revealed many molecular components of the AIS, their functions and interactions, and above all the importance of the AIS in the nervous system. Much less is known about the second ultrastructural hallmark of the AIS: the bundling of its microtubules into tight groups (“fascicles”) by electron-dense cross-bridges (Palay et al., 1968; Peters et al., 1968). The function of microtubule fascicles and the molecular basis of their formation were unknown until the discovery of the microtubule-associated protein tripartite motif containing 46 (TRIM46) at the AIS (van Beuningen et al., 2015).

TRIM46 is the reported driver of AIS microtubule fasciculation. Microtubules fail to form fascicles after TRIM46 knockdown in cultured neurons, while TRIM46 expression in HeLa cells is sufficient to induce microtubule fasciculation (van Beuningen et al., 2015; Harterink et al., 2019). TRIM46 knockdown is also reported to impair axon specification and AIS formation, both in cultured neurons and in neonatal mice after *in utero* electroporation with TRIM46 shRNA (van Beuningen et al., 2015). These findings strongly supported a key role for TRIM46 in axon specification and AIS formation and structure and provided a conceptually satisfying explanation for the role of bundled AIS microtubules. However, these conclusions were based mostly on *in vitro* studies and acute knockdown of TRIM46. Here, we examined TRIM46 knockout mice to determine the consequences of TRIM46 loss on the nervous system and to define the relationship between TRIM46, AnkG, and AIS formation; AIS formation follows axon specification. We found that loss of TRIM46 *in vivo* has minimal consequences for mouse health and behavior and, contrary to prior results, does not prevent AIS formation or axon specification. We also describe the presence of TRIM46 at proximal nodes of Ranvier throughout the nervous system. In all, our findings suggest that the relatively severe consequences of TRIM46 loss reported in other systems do not replicate *in vivo*. These results underscore the need to search for mechanisms controlling microtubule polarity and function at the AIS. Our study also demonstrates the importance of validating conclusions derived from highly reduced and simplified model systems (e.g. primary neuronal culture) in more complex animal models.

## MATERIALS AND METHODS

### Animals

C57BL/6NJ-*Trim46^em1(IMPC)J^*/Mmjax mice (RRID:MMRRC_066813-JAX) were obtained from the Mutant Mouse Resource and Research Center (MMRRC) at The Jackson Laboratory, an NIH-funded strain repository. Mice were genotyped by PCR of genomic DNA extracted from ear punches with the following primers: forward: 5’ cagatcgcttggaccggctccttaa 3’, reverse: 5’ gctgcaattacagggaaacaaatgtgattctct 3’. To sequence the excision site, genomic DNA was amplified with the following primers: forward: 5’ gcagtaggtagcttgtgc 3’, reverse: 5’ gccaggctctttgttagc 3’. The purified product was then Sanger sequenced by Genewiz with the forward genotyping primer. *ChAT-Cre;Ank3^F/F^* and *Adv-Cre;Ank3^F/F^* mice were generated as previously described (Teliska et al., 2022). Timed pregnant Sprague-Dawley rats were obtained from Charles River Laboratories. All experiments complied with the National Institutes of Health’s Guide for the Care and Use of Laboratory Animals and were approved by the Baylor College of Medicine Institutional Animal Care and Use Committee.

### Antibodies

The following primary antibodies were used for immunofluorescence: mouse monoclonal antibodies against AnkG (UC Davis/NIH NeuroMab Facility clone N106/65, RRID:AB_2877525, 1:200 and clone N106/36, RRID:AB_2877524, 1:150-1:200), FOXP2 (Atlas Antibodies Cat# AMAb91361, RRID:AB_2716651, 1:500), Myc (MBL International Cat# M192-3, RRID:AB_11160947, 1:2000), NeuN (Sigma-Aldrich Cat# MAB377, RRID:AB_2298772, 1:200-1:500), and Reelin (Millipore Cat# MAB5364, RRID:AB_2179313, 1:500), rabbit monoclonal antibodies against NF186 (Cell Signaling Technology Cat# 15034 (also 15034S), RRID:AB_2773024, 1:1000) and TRIM46 (#1 in text: Abcam Cat# ab307967, RRID:AB_3096363, 1:200-1:1000, #2 in text: Cell Signaling Technology Cat# 92574, RRID:AB_3096364, 1:200), rabbit polyclonal antibodies against βIV-spectrin (M.N. Rasband, Baylor College of Medicine, Texas, USA Cat# bIV SD, RRID:AB_2315634, 1:500), Caspr (M.N. Rasband, Baylor College of Medicine, Texas, USA Cat# Anti-Caspr, RRID:AB_2572297, 1:250-1:500), CUX1 (Santa Cruz Biotechnology Cat# sc-13024, RRID:AB_2261231, 1:100), Na_v_1.6 (Alomone Labs Cat# ASC-009, RRID:AB_2040202, 1:500), and NDEL1 (Thermo Fisher Scientific Cat# PA5-53669, RRID:AB_2644530, 1:200), guinea pig monoclonal antibody against TRIM46 (Synaptic Systems Cat# 377 308, RRID:AB_2924929, 1:1000), guinea pig polyclonal antibody against TRIM46 (Synaptic Systems Cat# 377 005, RRID:AB_2721101, 1:500-1:1000), rat monoclonal antibody against CTIP2 (BioLegend Cat# 650602 (also 650601), RRID:AB_10915967, 1:200), and chicken polyclonal antibody against MAP2 (EnCor Biotechnology Cat# CPCA-MAP2, RRID:AB_2138173, 1:1000).

The following primary antibodies were used for immunoblotting: mouse monoclonal antibodies against TuJ1 (BioLegend Cat# 801202 (also 801201, 801213), RRID:AB_10063408, 1:800 and Santa Cruz Biotechnology Cat# sc-58888, RRID:AB_1119489, 1:500), rabbit monoclonal antibodies against TRIM46 (#1 in text: Abcam Cat# ab307967, RRID:AB_3096363, 1:500, #2 in text: Cell Signaling Technology Cat# 92574, RRID:AB_3096364, 1:1000), guinea pig monoclonal antibody against TRIM46 (Synaptic Systems Cat# 377 308, RRID:AB_2924929, 1:500), and guinea pig polyclonal antibody against TRIM46 (Synaptic Systems Cat# 377 005, RRID:AB_2721101, 1:500).

Secondary antibodies were purchased from Thermo Fisher Scientific, Jackson ImmunoResearch Laboratories, and Aberrior.

### Epitope mapping

To generate full-length, truncated, and mutant TRIM46 constructs, regions of *Trim46* were amplified with PCR from full length mouse *Trim46L* obtained from the GFP-Trim46 plasmid previously described in (van Beuningen et al., 2015) (gift from Drs. Casper Hoogenraad & Thomas Schwarz (Addgene plasmid # 176402; http://n2t.net/addgene:176402; RRID:Addgene_176402)). A point mutation in this plasmid (A681T) was carried through to our TRIM46L constructs. The Ensembl sequence and exon annotation for mouse *Trim46* (ENSMUSG00000042766) and was used to design primers to make the appropriate truncations. The primers were designed to add a C-terminal Myc tag to the amplicons. SnapGene software was used for plasmid/primer design. To generate the TRIM46S and TRIM46LΔ constructs, which contain sequences not found in TRIM46L (for TRIM46S: the first and last exon, for TRIM46LΔ: 14 frameshifted residues of exon 4 after exon 2), the primers were designed to append the necessary sequences at the appropriate loci. Amplicons were inserted into a pAAV-CMV vector using In-Fusion cloning (Takara Bio). Plasmids were verified with Sanger sequencing (Genewiz) or whole plasmid sequencing (Plasmidsaurus). The ranges of each TRIM46L construct within full length 759aa TRIM46L (UniProt Q7TNM2) are as follows: TRIM46L (1-759aa), TRIM46L E1-3 (1-223aa), TRIM46L E1-2 (1-108aa), TRIM46LΔ (1-108aa + GSDVSRPQGRGDPLL), TRIM46L E3-11 (109-759aa). The ranges of each TRIM46S construct within full length 541aa TRIM46S (UniProt Q3TC52) are as follows: TRIM46S (1-541aa), TRIM46S E1-2 (1-95aa), TRIM46S E3-11 (96-541aa).

Constructs were transfected into COS7 cells plated on glass coverslips. 48 hours later, cells were fixed with 4% paraformaldehyde (PFA), pH 7.2, for 10 minutes at room temperature. Cells were washed with PBS or PB and then blocked with 0.1 M PB containing 0.3% Triton X-100 and 10% normal goat serum (PBTGS) for at least 30 minutes before incubation in primary antibody diluted in PBTGS for 1 hour-overnight. Cells were washed with PBTGS, incubated with secondary antibody diluted in PBTGS for 45 minutes-1 hour, washed with PBTGS and PB, and mounted. All staining steps were performed at room temperature.

### Behavioral testing

All behavioral tests were performed and analyzed by the same person, who handled the mice 1-2 times per day for at least 3 days before beginning testing. Mice were allowed to habituate to the testing room for at least 1 hour prior to each test. Behavioral apparatuses were cleaned with ethanol between animals. No two behavioral tests were done on the same day. The experimenter was blinded to genotype during testing and analysis.

### Open field test and elevated plus maze test

The open field test was performed for 10 minutes in an open-top acrylic box with a light bottom and dark sides (40 × 40 × 20 cm). The elevated plus maze test was performed for 10 minutes. The arms of the maze were 30 × 5 cm and the maze was raised 40 cm off the floor. Mouse movement was tracked with a camera and analyzed with ANY-maze software.

### Rotarod

Mice underwent three trials on the rotarod (Ugo Basile) per day for three consecutive days accelerating from 4 rpm to 40 rpm over 5 minutes. Prior to each trial on the first day, mice were acclimated to the rotarod for 1 minute at 4 rpm. Latency to fall was recorded for each trial. If a mouse ceased to walk on the rotarod and instead held on for two full rotations, it was counted as a fall. If a mouse fell in the first 10 seconds, it was put back on the rotarod. Mice rested for at least 30 minutes between trials.

### Wire hang

Mice were placed on a wire mesh grid and then inverted 36 cm above a cage filled with bedding nestlets for cushioning. The inversion was done slowly to allow the mice to grasp the wire with all four paws as they were turned. Mice were acclimated to the wire hang with the first trial and then their latency to fall was recorded for three subsequent trials with a cut-off time of 3 minutes. Mice rested for at least 30 minutes between trials.

### Footprint analysis

The hindfeet of mice were painted with washable, non-toxic paint and then mice were placed in the entrance of a 45 cm clear plexiglass tunnel that ended in a dark chamber. Bright overhead light shone on the tunnel to encourage mice to walk through it to the chamber. White paper was placed in the bottom of the tunnel to collect the footprints. Mice walked through the tunnel multiple times to gather enough footprints to take at least nine measures of stride, stance, and sway. Stride was measured as the distance from one left footprint to the next, stance as the diagonal distance from one left footprint to the next right footprint, and sway as the straight distance between adjacent left and right footprints. Measurements were done in FIJI/ImageJ.

### Tissue immunofluorescence

Adult mice were euthanized by isoflurane overdose followed by decapitation. Postnatal day 0 mice were euthanized by ice anesthesia followed by decapitation. Tissues were dissected and fixed in 4% paraformaldehyde, pH 7.2, on ice for 1 hour (brain) or 30-35 minutes (spinal cord, nerves, roots, and DRGs). The DRGs in Fig. 10E were fixed 1 hour in 2% PFA, pH 7.2, on ice.

Tissues were then cryoprotected at 4 °C in 20 or 30% sucrose in 0.1 M phosphate buffer (PB) or moved to PB for floating immunostaining (Fig. 10A, D). For floating staining, all steps were performed on a rocker at room temperature. Tissues were permeabilized in 2% Triton X-100 in PB for 30 minutes, then blocked with PBTGS for 1 hour. They were then incubated with primary antibody diluted in PBTGS overnight, washed with PBTGS, incubated with secondary antibody in PBTGS for 2 hours, and finally washed with PBTGS followed by PB before mounting.

For slide-mounted staining, tissues were embedded in Tissue-Tek OCT (Sakura Finetek 4583) and cryosectioned with a Thermo Fisher Scientific Cryostar NX70 or Thermo Fisher Scientific HM525 NX cryostat at 25 μm (brain and spinal cord) or 14 μm (dorsal roots and DRGs). Sections were collected onto glass coverslips coated with 1% bovine gelatin, dried at room temperature, washed with PB, and blocked for 1 hour in PBTGS. Sections were then incubated overnight in primary antibody diluted in PBTGS, washed with PBTGS, incubated in secondary antibody dilution in PBTGS for 1-2 hours, washed with PBTGS and PB, dried, and mounted. In some cases, sections were treated with TrueBlack Lipofuscin Autofluorescence Quencher (Biotium). All staining was done at room temperature. The variations in fixation and staining conditions in these methods reflect variations between experiments, not within an experiment. Conditions were identical between groups within each experiment.

### Immunoblotting

Adult mice were euthanized by isoflurane overdose followed by decapitation. Brains were quickly dissected and immediately homogenized or flash frozen on dry ice and stored at -80 °C for later homogenization. Brains were homogenized on ice in a glass Dounce Homogenizer with ice-cold homogenization buffer containing 0.32 M sucrose, 5 mM sodium phosphate buffer, pH 7.2, 1 mM NaF, 1 mM Na_3_VO_4_, 0.5 mM PMSF, and protease inhibitors. Homogenates were centrifuged at 700 × g for 10 minutes at 4 °C to remove nuclei and cell debris. The supernatant was then centrifuged at 4 °C either at 20,800 × g for 1 hour or 27,200 × g for 1.5 hours.

Centrifugation speed/time was identical between samples on the same blot. The pellet was then resuspended in homogenization buffer. Protein concentration was measured with a Bradford assay (Bio-Rad) and samples were prepared for SDS-PAGE by dilution in SDS sample buffer and heating at 95 °C for 5 minutes. 20 or 30 μg was used per sample; this amount was kept identical between samples run on the same gel and is specified in the figure legend.

Samples were resolved on 8% polyacrylamide gels with electrophoresis and transferred onto nitrocellulose membranes using a Bio-Rad Trans-Blot Turbo transfer system. Membranes were washed with PBS and blocked with Blotto (5% dry skim milk and 0.05% Tween-20 in TBS) at room temperature for at least 40 minutes. Primary antibodies were diluted in Blotto and incubated for 1 hour at room temperature or overnight at 4 °C. HRP-conjugated secondary antibodies were diluted in Blotto and incubated for 30 minutes-1 hour at room temperature. Membranes were washed with Blotto after primary antibody incubation and with PBS containing 0.1% Tween-20 after secondary antibody incubation. Membranes were developed with SuperSignal West Pico PLUS (Thermo Scientific cat# 34580) or SuperSignal West Femto Maximum Sensitivity Substrate (Thermo Scientific cat# 34095) and imaged with a Licor Odyssey FC Imaging System.

### Neuron culture, CRISPR, and detergent extraction

Timed pregnant Sprague-Dawley rats from Charles River Laboratories were euthanized for embryo collection at E18. Hippocampi were dissected and dissociated. Neurons were plated on glass coverslips coated with Poly-D-Lysine (Sigma cat# P7886) and laminin (Life Technologies cat# 23017015) at a density of ∼1.25 × 10^4^ cells/cm^2^. Cortices were dissected in the same protocol and plated fresh or cryopreserved for culture as described previously (Ishizuka and Bramham, 2020). The cryopreserved neurons were plated at a density of ∼5 × 10^4^ cells/cm^2^.

Hippocampal and cortical neurons were maintained in Neurobasal medium (Life Technologies cat# 21103049) containing 1% Glutamax (Life Technologies cat# 35050061), 1% penicillin-streptomycin (Life Technologies cat# 15140122), and 2% B-27 supplement (Life Technologies cat# 17504044) in an incubator with 5% CO2 at 37 °C. To knock out AnkG, an adeno-associated viral (AAV) vector- and CRISPR-based strategy was employed, as previously described (Ogawa et al., 2023; Zhang et al., 2023). Small-scale AAV cell lysates were generated by triple-transfection with AAV plasmid, helper plasmid (Agilent Technologies, Cat # 240071), and serotype PHP.S plasmid (a gift from Dr. Viviana Gradinaru, Addgene plasmids #103006) with PEI Max (Polysciences, Cat # 24765) in HEK293T cells, as previously described (Ogawa et al., 2023; Zhang et al., 2023). The triple guide RNA knockout construct for rat *Ank3* has been described (Zhang et al., 2023). Triple guide RNAs for mouse and rat *Trim46* were designed: 5’ ctgtaagacatgtcaacgac 3’, 5’ aaacctgaccctagagcgag 3’, 5’ accggctccttaagtcaggt 3’. Cultured hippocampal neurons were co-infected with AAVs expressing the *Ank3* or *Trim46* knockout gRNAs and Cas9 3-5 hour after plating. Cultured cortical neurons were infected with AAVs 1-5 days after plating. A full media change was performed 2 days after AAV infection. Thereafter, half of the media was removed and replaced with fresh media every 5 days. Cultured neurons were fixed at day *in vitro* (DIV) 17-21 in 4% PFA, pH 7.2, for 15 min at 4 °C and washed three times in 1X PBS. Fixed neurons were permeabilized and blocked with PBTGS for 1 hour at room temperature. Neurons were then incubated in primary antibodies diluted in PBTGS for 1 hour to overnight at room temperature. After incubation with primary antibodies, neurons were washed 3 times with PBTGS. Secondary antibodies were diluted in PBTGS and added to neurons for 1 hour at room temperature. Neurons were then washed with PBTGS and PB and mounted. For detergent extraction studies, neurons were incubated with ice-cold 1X PBS containing 0.5% Triton X-100 at 4 °C for 30 minutes before fixation and immunostaining. Neurons were then washed with ice-cold 1X PBS 3 times at 4 °C and fixed.

### Image analyses

Widefield images were collected on a Nikon Eclipse Ni-E microscope or a Zeiss AxioImager with AxioCam. The Zeiss microscope was fitted with an ApoTome which was sometimes used for optical sectioning. Some images were taken as z-stacks and are displayed as maximum intensity projections (when taken on the Zeiss microscope) or Extended Depth of Focus images (when taken on the Nikon microscope and processed with Nikon Elements software). Confocal z-stacks were collected on a Nikon Eclipse Ti2 microscope fitted with an Abberior STEDYCON system and are displayed as maximum intensity projections. FIJI/ImageJ was used to crop and merge images, make linear contrast adjustments, and, in some cases, perform rolling ball background subtraction. For Fig. 5, two serial sections were stained, each for two of the four layer markers shown. Images were then collected from each section in the same region, and the images were aligned and overlaid based on pial surface in Adobe Photoshop.

Image quantification was done in FIJI/Image using the Cell Counter plugin (Dr. Kurt De Vos), NeuronJ plugin (Meijering et al., 2004), Measure ROIs macro set (Dr. Christophe Leterrier), and Blind Analysis Tools plugin (Drs. Astha Jaiswal and Holger Lorenz). For intensity and length measurements of AnkG staining, images were processed with rolling ball background subtraction and a Gaussian blur. The start and end of the AnkG+ region was taken at the first and last point where the AnkG intensity was above 35% of its maximum. For length measurements of TRIM46 staining, a Gaussian blur was used and the start and end were taken at the first and last point where TRIM46 intensity was above 20% of its maximum. For quantification of the percentage of neurons *in vivo* with an AIS, a mask marking neurons was generated based on NeuN staining. Marked neurons with and without AIS (judged by AnkG staining) were then counted. Marked neurons that could not be assessed (i.e. because they were at the edge of the field of view) were excluded. For quantification of the percentage of AIS with βIV-spectrin, NF186, NDEL1, and Na_v_1.6, AIS were first marked based on AnkG staining, without looking at staining for the other proteins, then the number of marked AIS with and without the other proteins was counted. For quantification of cultured neurons, neurons were first marked for assessment based on MAP2 staining, without looking at other channels.

Marked neurons were then classified as having AIS containing AnkG, TRIM46, both, or neither. Neurons that could not be assessed (i.e. those growing in clumps) were excluded. All quantifications were done blinded, except for measurement of TRIM46 length because the difference between conditions was obvious.

### Statistical Analyses

Data were analyzed using GraphPad Prism and Microsoft Excel. Statistical tests, sample sizes, and error bars are described in figure legends. Data distribution was assumed to be normal.

## RESULTS

### TRIM46 knockout mice are valid and viable

To investigate the function of TRIM46 *in vivo*, we examined *Trim46* constitutive knockout (*Trim46^-/-^*) mice. Mice were generated by CRISPR-mediated deletion of 435 bp encompassing the entire third exon of *Trim46*, resulting in a frameshift mutation and premature stop codon early in the fourth exon (Fig. 1A). We confirmed the deletion by PCR and Sanger sequencing of genomic DNA (Fig. 1C, E). To test the validity of the knockout, we performed immunostaining and immunoblotting of adult *Trim46^-/-^* brain with four different TRIM46 antibodies (Fig. 1F-M). Immunostaining of wildtype cortex revealed strong AIS TRIM46 labeling, while no immunoreactivity was detected in *Trim46^-/-^* cortex with any of the antibodies (Fig. 1F-I). Immunoblotting of brain homogenate using three of the four antibodies similarly suggested complete loss of TRIM46 protein (Fig. 1J-L), while the fourth antibody did not work for immunoblotting in our hands (Fig. 1M).

**Fig. 1.**
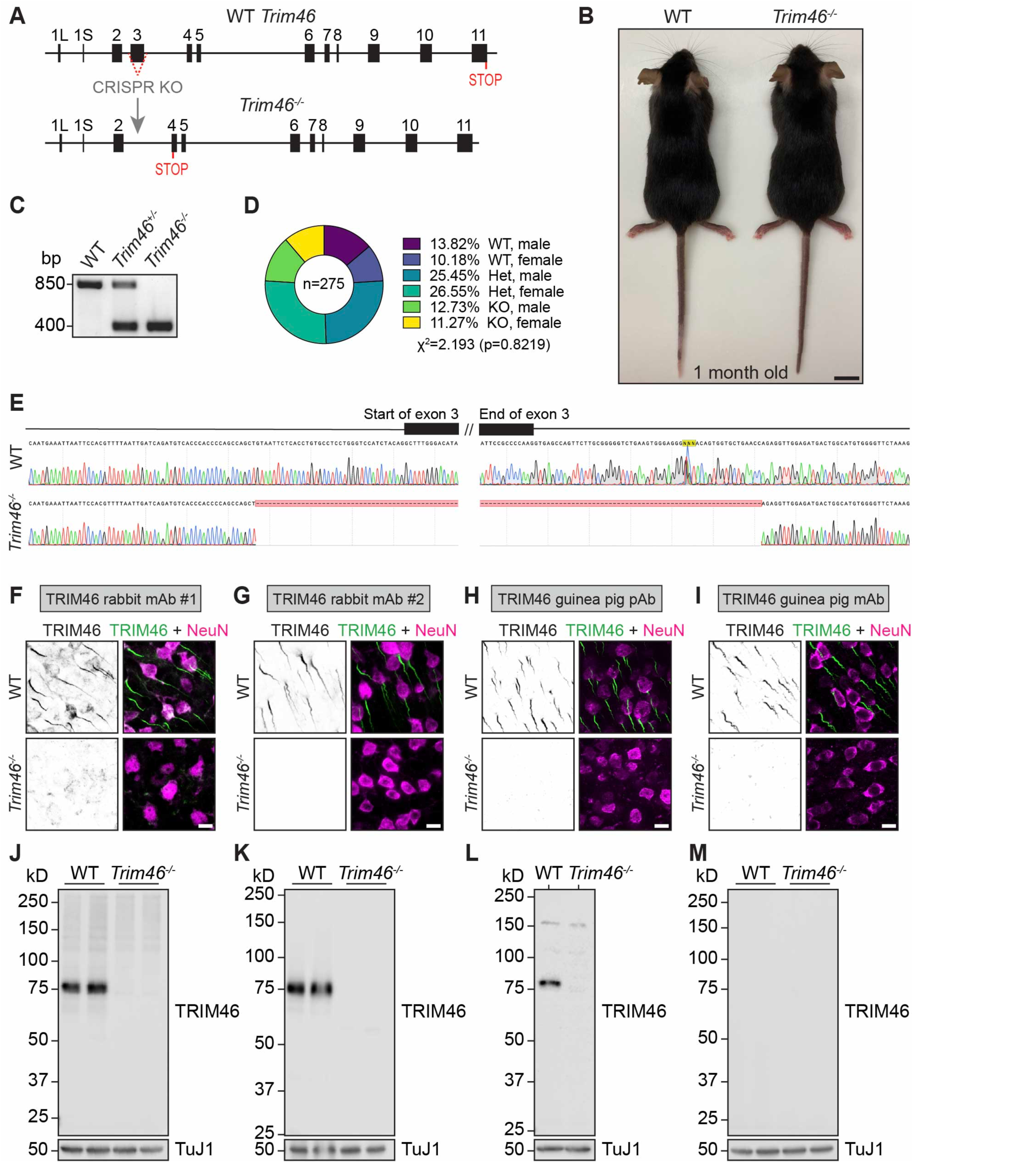
Validation of TRIM46 knockout mice. (A) Design of knockout allele. (B) Images of 1-month old wildtype and *Trim46^-/-^* mice. (C) PCR of genomic DNA. (D) Pie chart of genotypes obtained from crossing *Trim46^+/-^* mice (n=275). (E) Sanger sequencing of genomic DNA at the start and end of exon 3. (F-M) Validation of TRIM46 knockout by immunostaining (F-I) and immunoblotting (J-M) using four different TRIM46 antibodies: two monoclonal rabbit antibodies (#1: F, J; #2: G, K), a polyclonal guinea pig antibody (H, L), and a monoclonal guinea pig antibody (I, M). (F-I) Immunostaining of TRIM46 (green) and NeuN (magenta, shown only in the merged images) in adult cortex. (J-M) Immunoblotting of 20 μg (J, K, M) or 30 μg (L) crude synaptosomal fraction of adult brain. Scale bars: (B) 1 cm, (F-I) 10 μm.

*Trim46^-/-^* mice are viable and appear grossly normal (Fig. 1B). They do not exhibit perinatal lethality, as heterozygote crosses produce genotypes consistent with Mendelian ratios when genotyped at postnatal day 15 or older (Fig. 1D, n=275 mice from 39 litters). We aged some *Trim46^-/-^* mice beyond 18 months of age, though we did not formally quantify lifespan. Given that TRIM46 is reported to be required for AIS formation (van Beuningen et al., 2015) and that AnkG knockout mice, which cannot form AIS, die at birth (Jenkins et al., 2013, 2015; Ho et al., 2014), the viability of *Trim46^-/-^* mice underscored the need to rigorously validate the knockout. Therefore, we next mapped the epitopes of the antibodies used in Figure 1 to address the possibility that *Trim46^-/-^* mice express a mutant TRIM46 fragment undetectable by our antibodies.

The *Trim46* gene produces two known protein isoforms, TRIM46L and TRIM46S. They differ in their first exon, the inclusion of exon 10 (which TRIM46L includes and TRIM46S skips), and their last exon. *Trim46S* is primarily transcribed before neuronal differentiation and it produces an unstable peptide undetectable in brain (Vuong et al., 2022). Though this suggests that TRIM46L is the only functional TRIM46 isoform in neurons, for the sake of thoroughness we considered both isoforms in our experiments. As the isoforms are identical from exons two through nine, the deletion of exon three in the knockout allele should cause an identical frameshift in both, appending a short (14 aa) mutated sequence to exon two followed by a premature stop codon (Fig. 2A, red exon labeled “Δ”). We generated plasmids to express this hypothetical mutant as well as various N- and C-terminal truncations of both isoforms and transfected them into COS-7 cells (schematized in Fig. 2A with representative images in Fig. 2B). All plasmids included a C-terminal Myc tag.

**Fig. 2.**
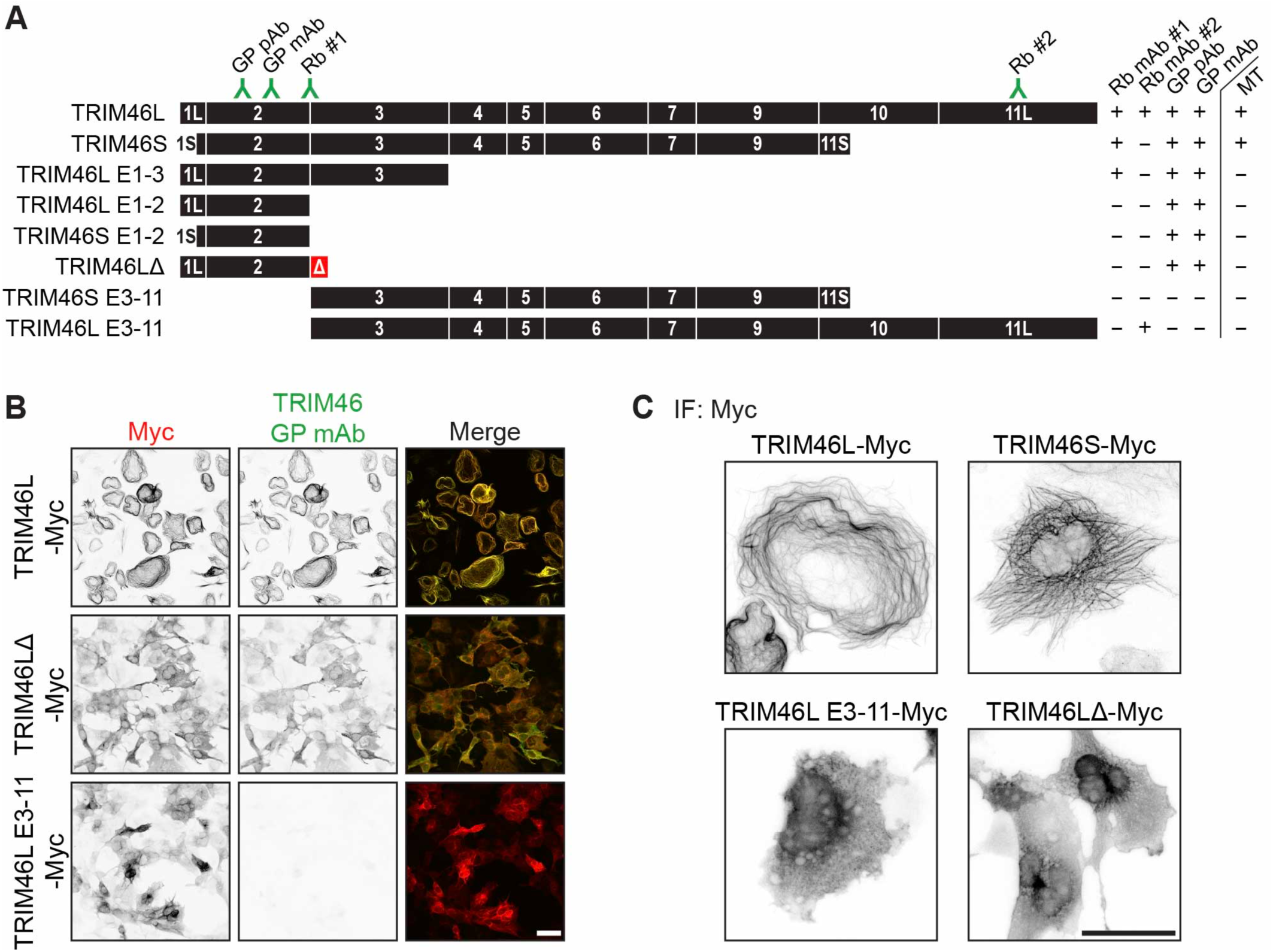
Epitope mapping of TRIM46 antibodies. (A-B) Epitope mapping of the four TRIM46 antibodies used in Figure 1, summarized in (A) with representative images in (B) (red: Myc tag on TRIM46 construct, green: TRIM46 antibody). (A) Also indicates microtubule localization of each construct. (C) Immunostaining showing subcellular distribution of various TRIM46 constructs. Scale bars: 50 μm.

We found that rabbit antibody #1 (Fig. 1F, J) labels a truncation that goes through exon three but not the ones that end at exon two or start at exon three. Therefore, its epitope bridges exons two and three, and it could not detect a hypothetical mutant lacking exon 3. Rabbit antibody #2 (Fig. 1G, K) labeled a C-terminal fragment of TRIM46L but not TRIM46S. Correspondence with the supplier revealed that it was generated against residues in the last exon of TRIM46L, which is not found in TRIM46S. Therefore, if the knockout allele produces a C-terminal TRIM46L fragment with an alternative start codon past the deleted exon 3, it would be detected by this antibody. The two guinea pig antibodies (Fig. 1H, I, L, M) labeled N-terminal fragments of both isoforms that end after exon two as well as the hypothetical mutant form of TRIM46L (TRIM46LΔ). Neither antibody labeled the N-terminal truncation that starts at exon three, indicating that their epitopes are contained entirely within the region of the protein that would be retained in the hypothetical N-terminal mutant. Together, the epitopes of these antibodies span N- and C-terminal regions of TRIM46, so the absence of immunostaining in *Trim46^-/-^* brain (Fig. 1F-I) indicates that there is no mutant TRIM46 protein present at the AIS. In summary, the results shown in Figures 1 and 2 demonstrate that *Trim46^-/-^* mice are a valid and viable *in vivo* model of TRIM46 knockout.

The epitope mapping experiment afforded us the opportunity to examine the subcellular localization of each of the truncates in transfected cells. All truncates had a diffuse cytosolic distribution, with only full length TRIM46L and TRIM46S showing extensive microtubule localization (Fig. 2A, “MT” column, representative images in Fig. 2C). This replicates the prior finding that the COS box in exons 6-7, a common motif among microtubule-binding members of the TRIM superfamily (Short and Cox, 2006), is required for TRIM46 to associate with microtubules (van Beuningen et al., 2015).

### TRIM46 is not required for axon specification and AIS formation in vivo

Because multiple prior studies report a role for TRIM46 in AIS formation *in vitro*, we first investigated whether AIS formation is impaired in *Trim46^-/-^* mice. Immunostaining for AnkG revealed abundant AIS in adult *Trim46^-/-^* brain (Fig. 3A). Since AIS assembly follows axon specification these results also show appropriate development of axon identity. Using NeuN as a neuronal marker, we found no reduction in the percentage of neurons with an AIS, the fluorescence intensity of AnkG, or the length of the AIS in adult *Trim46^-/-^* cortex (Fig. 3B-D). These findings indicate that, despite prior reports in cultured neurons, TRIM46 is not required for AIS formation or axon specification *in vivo*.

**Fig. 3.**
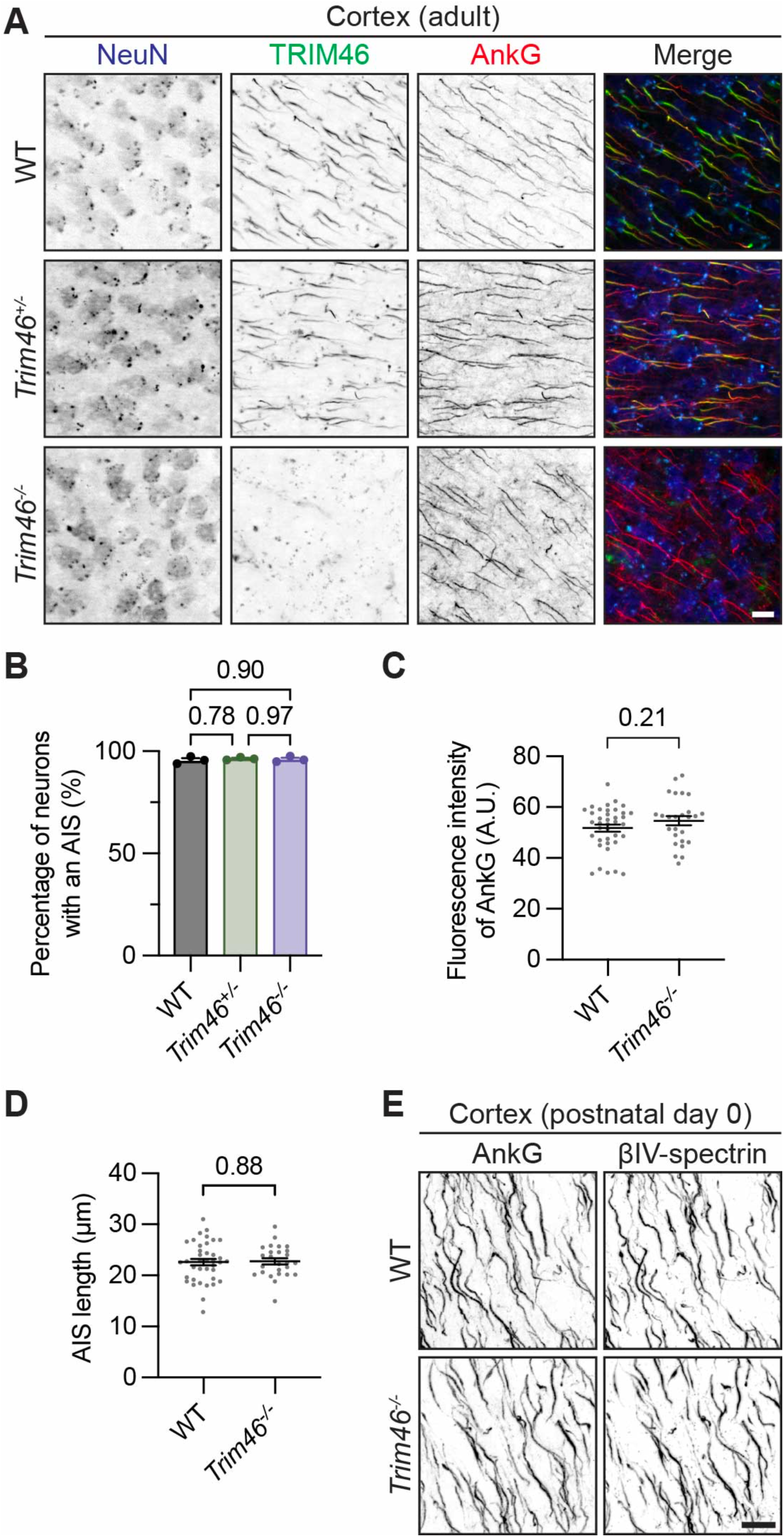
TRIM46 is not required for AIS formation *in vivo*. (A) Immunostaining of NeuN (blue), TRIM46 (green), and AnkG (red) in adult cortex. (B-D) Quantification of AIS in adult cortex of (B) percentage of neurons with an AnkG+ AIS (n=3 mice, N=211-422 neurons per mouse), (C) fluorescence intensity of AnkG at the AIS, and (D) AIS length (based on AnkG staining) (N=38 WT and 26 *Trim46^-/-^* AIS from n=1 mouse for C-D). (B) was analyzed with one-way ANOVA (p=0.7886) with Tukey’s multiple comparisons test (adjusted p-values shown on graph). (C-D) were analyzed with unpaired t-tests. (E) Immunostaining of AnkG and βIV-spectrin in cortex at postnatal day 0 (n=2). Error bars represent SEM. Scale bars: 10 μm.

Because of the departure of these findings from prior work, we questioned whether our results capture the full scope of TRIM46’s necessity for AIS formation. We considered the possibility that, though *Trim46^-/-^* mice have AIS by adulthood, TRIM46 knockout could cause a transient delay in AIS formation during development. This seemed unlikely, as mice lacking AIS are known to die at birth (Jenkins et al., 2013, 2015; Ho et al., 2014). Nevertheless, we tested this possibility by immunostaining brain from wildtype and *Trim46^-/-^* mice for AnkG and βIV-spectrin at postnatal day 0 (P0). We found that *Trim46^-/-^* brain has abundant AIS by P0 (Fig. 3E). Most AIS formation in the cerebral cortex begins before birth and finishes around P1 (Galiano et al., 2012), so this experiment does not rule out the possibility of a prenatal delay in *Trim46^-/-^* mice; however, these data show that if such a delay occurs at all, it is brief and resolves by birth.

AnkG is the master organizer of the AIS, but it is far from the only important AIS protein. To further examine the effect of TRIM46 knockout on AIS formation, we immunostained adult cortex for four additional AIS proteins: βIV-spectrin, Neurofascin-186 (NF186), Nuclear distribution element-like 1 (NDEL1), and the voltage gated sodium channel Na_v_1.6. We found these proteins at AIS throughout *Trim46^-/-^* cortex (Fig. 4A-D). Using AnkG as an AIS marker, we quantified the percentage of AIS with each protein and found no significant differences between *Trim46^-/-^* and control mice (Fig. 4E). Together, these data suggest that AIS formation is not impaired in *Trim46^-/-^* mice.

**Fig. 4.**
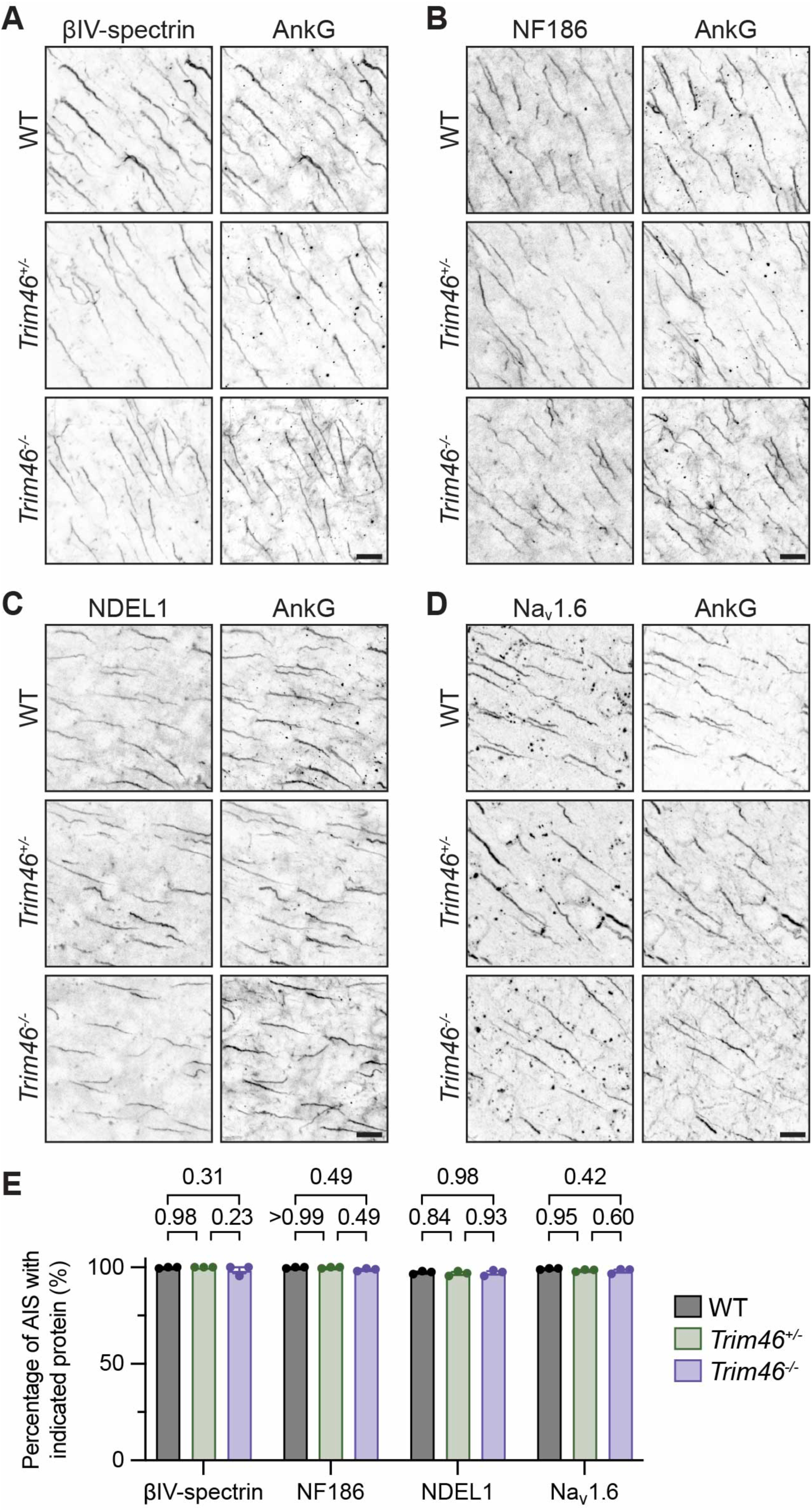
TRIM46 is not required for the localization of major AIS proteins *in vivo*. (A-D) Immunostaining and (E) corresponding quantification in adult cortex for AIS proteins (A) βIV-spectrin, (B) Neurofascin186, (C) NDEL1, and (D) Nav1.6, all co-labeled with AnkG (n=3 mice, N=143-200 AIS per mouse). (E) Percentage of AIS in adult cortex with each protein. Analyzed with two-way ANOVA (p_genotype_=0.1319) with Tukey’s multiple comparisons test (adjusted p-valued shown on graph). Error bars represent SEM. Scale bars: 10 μm.

### TRIM46 knockout mice have intact cortical lamination

Previous studies reported impaired cortical neuron migration after *in utero* electroporation with TRIM46 shRNA (van Beuningen et al., 2015). To determine if TRIM46 knockout impairs neuronal migration, we immunostained adult *Trim46^-/-^* cortex for cortical layer markers Reelin (layer I), CUX1 (layers II-IV), CTIP2 (layers V-VI), FOXP2 (primarily layer VI). The distribution of these markers was similar between *Trim46^-/-^* and wildtype cortex, indicating that cortical lamination is intact in *Trim46^-/-^* mice (Fig. 5). Though these data do not rule out the possibility that cortical neuron migration is delayed in developing *Trim46^-/-^* brain, they show that if such a delay does occur, it resolves by adulthood.

**Fig. 5.**
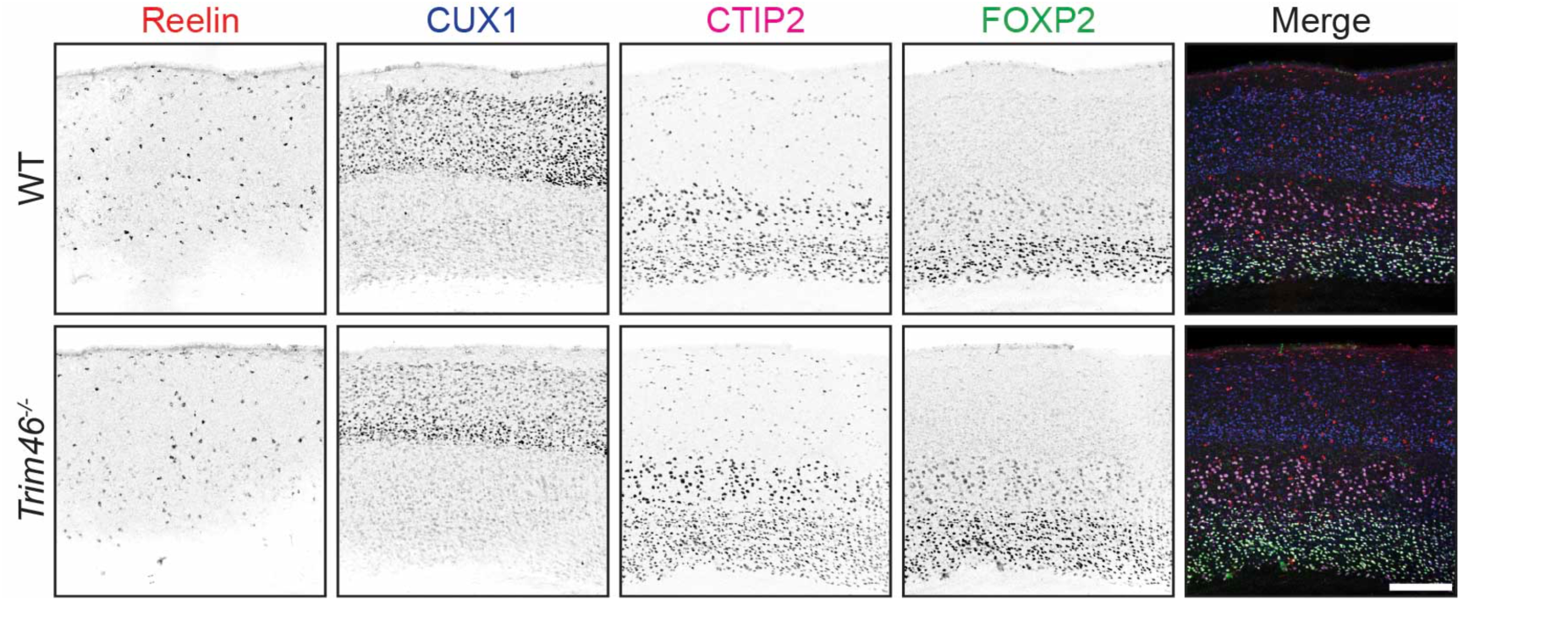
TRIM46 knockout mice have normal cortical lamination. Adult cortex immunostained for cortical layer markers Reelin (red), CUX1 (blue), CTIP2 (magenta), and FOXP2 (green). Scale bar: 200 μm.

### TRIM46 knockout mice exhibit mild motor deficits and normal anxiety-related behavior

To determine if TRIM46 knockout leads to behavioral changes, we performed a range of behavioral assays on adult *Trim46^-/-^* mice. We first tested motor function using rotarod, wire hang, and footprint analysis. On the rotarod, while wildtype mice improved over the course of nine trials, *Trim46^-/-^* mice of both sexes exhibited a decreased latency to fall at the start and little improvement over the testing period. Heterozygous males performed similarly to knockout males at first but improved to match the wildtype males by the end of the testing, while heterozygous females performed similarly to wildtype females from the start (Fig. 6A).

**Fig. 6.**
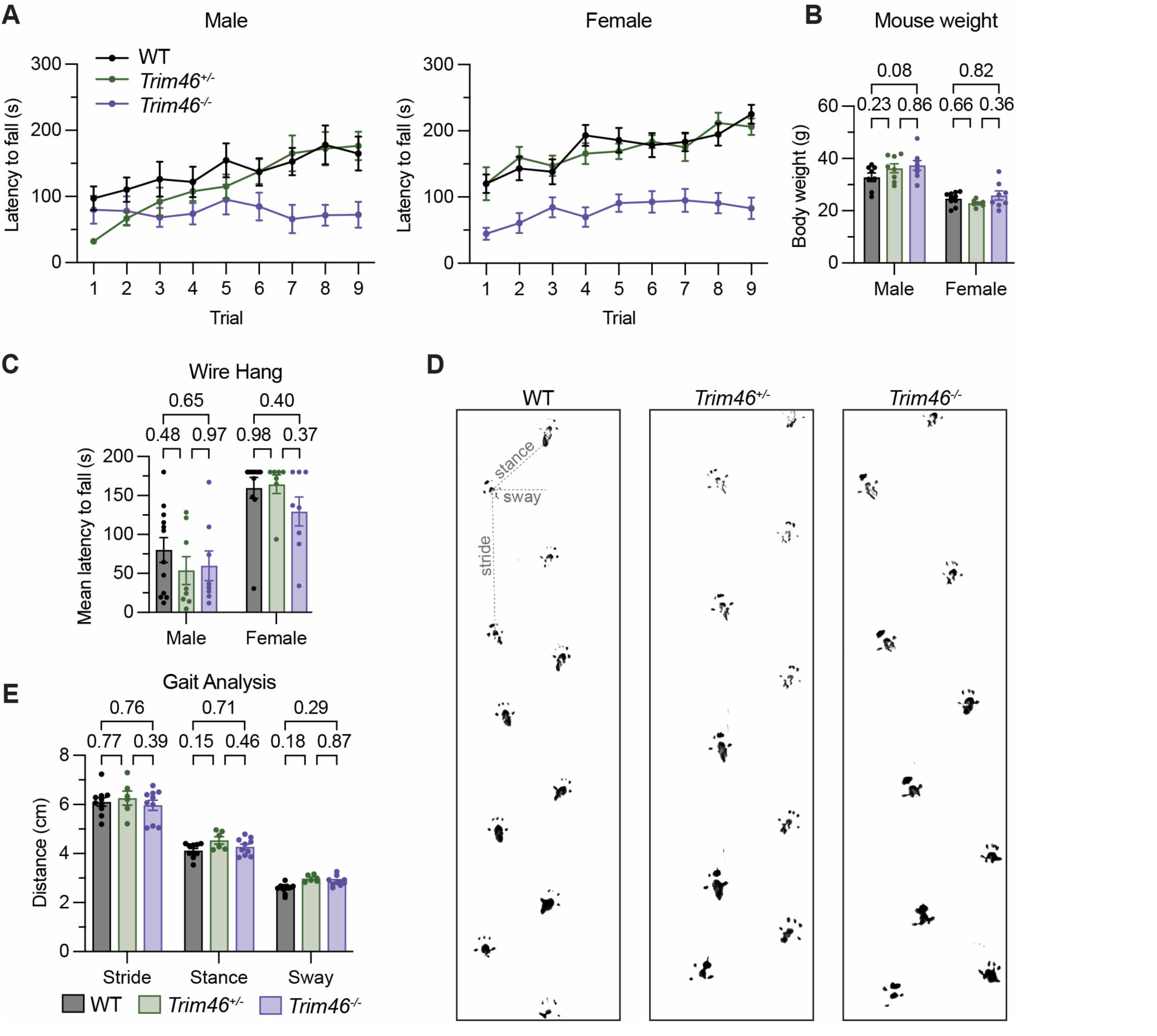
TRIM46 knockout mice exhibit a mild motor deficit. (A) Latency to fall on accelerating rotarod at 3-5 months of age (males, n=8; females, n=10 WT, 7 *Trim46^+/-^*, 8 *Trim46^-/-^*). The first point for *Trim46^+/-^* males lacks an error bar because the data were too close together to display one. Two-way ANOVA: for males, p_genotype_<0.0001, p_trial_=0.0001, p_genotype*trial_=0.0531; for females, p_genotype_<0.0001, p_trial_<0.0001, p_genotype*trial_=0.6068. (B) Body weight of the mice tested on rotarod and wire hang. (C) Average latency to fall on wire hang test at 3-5 months of age. (D, E) Gait analysis at 3-4 months of age. (B, C, E) were analyzed with two-way ANOVA (p-values: (B) p_genotype_=0.1367, (C) p_genotype_=0.3058, (E) p_genotype_=0.0466) with Tukey’s multiple comparisons test (adjusted p-values shown on graphs). Error bars represent SEM.

Analysis of the mice tested on rotarod confirmed that body weight did not differ significantly between genotypes and so could not explain the poor performance of *Trim46^-/-^* mice (Fig. 6B). On wire hang, no significant differences were observed in the mean latency to fall over three trials for either sex (Fig. 6C). For footprint analysis, two-way ANOVA indicated a significant effect of genotype as a source of variation (p=0.0466), but further analysis of stride, stance, and sway with Tukey’s multiple comparisons test did not identify significant differences for any measure (Fig. 6E). On qualitative assessment, the footprint tracks did not show obvious gait abnormalities in *Trim46^-/-^* mice (Fig. 6D).

To assess anxiety-like behavior and general locomotor activity, we used the open field test and the elevated plus maze, each with a 10 minute testing period (Fig. 7). Neither test revealed any significant differences in total distance traveled or time spent in the zone of the test field considered to be more anxiety-provoking (the inner zone for open field test and the open arm for elevated plus maze). Together, these behavioral data suggest that *Trim46^-/-^* mice exhibit mild motor impairment but otherwise normal behavior.

**Fig. 7.**
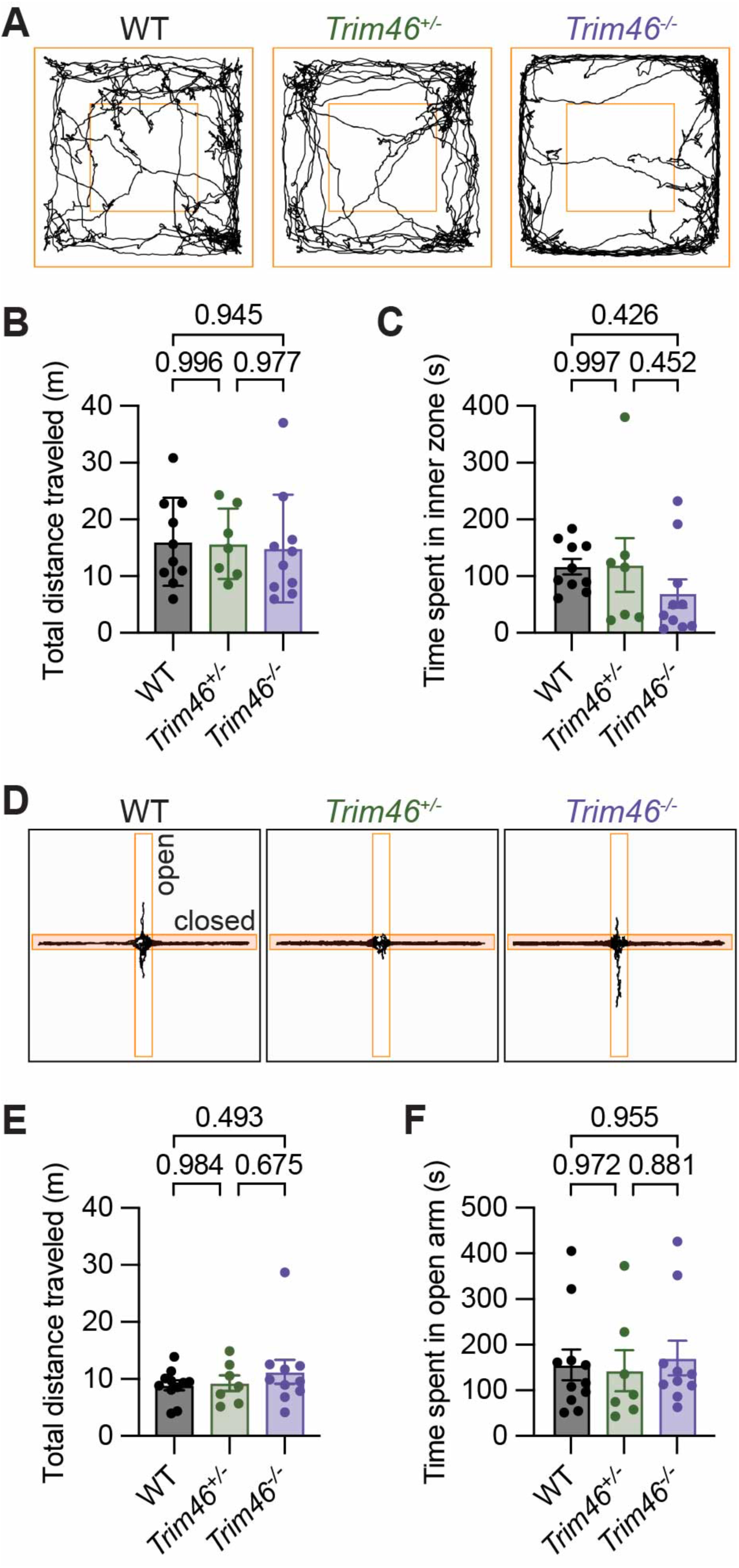
TRIM46 knockout mice do not exhibit anxiety-like phenotypes. (A-C) Open field test at 4-5 months of age: (A) representative traces, (B) total distance traveled, and (C) time spent in inner zone. (D-F) Elevated plus maze at 4-5 months of age: (D) representative traces, (E) total distance traveled, and (F) time spent in open arm. Analyzed with one-way ANOVA (p-values: (B) 0.9573, (C) 0.3594, (E) 0.4934, (F) 0.8891) with Tukey’s multiple comparisons test (adjusted p-values shown on graphs). Error bars represent SEM.

### TRIM46 localization depends on AnkG in vivo

After establishing that TRIM46 is not required for AnkG localization to the proximal axon *in vivo*, we next investigated the inverse relationship: the necessity of AnkG for TRIM46 localization. Prior work found that AnkG knockdown in cultured neurons induces a mislocalization of TRIM46: it accumulates in the cell body and/or extends into the distal axon (van Beuningen et al., 2015). To determine if this also occurs *in vivo*, we generated mice lacking AnkG in their spinal motor neurons by crossing *Ank3^F/F^* mice with *ChAT-Cre* mice (*ChAT-Cre;Ank3^F/F^*) as previously described (Teliska et al., 2022). We immunostained spinal cord from *ChAT-Cre;Ank3^F/F^* and control *Ank3^F/F^* mice and observed a striking extension of TRIM46 immunoreactivity much farther down the axon in AnkG-deficient neurons than in control neurons (Fig. 8A-B). This is consistent with the TRIM46 redistribution previously reported after AnkG knockdown in cultured neurons (van Beuningen et al., 2015). Quantification of the length of TRIM46-immunoreactive axon revealed that, on average, TRIM46 extends roughly four times farther in AnkG-deficient axons than in control axons (Fig. 8C). The TRIM46-positive region of *ChAT-Cre;Ank3^F/F^* axons is so long that in many cases we observed it either cross paths with other axons or stop bluntly instead of tapering off as it does in control neurons, suggesting that the true end was cut off during cryosectioning. Thus, our measurements may underestimate its length. These results show that TRIM46 does not require AnkG for axonal localization, but instead requires AnkG for its restriction to the AIS.

**Fig. 8.**
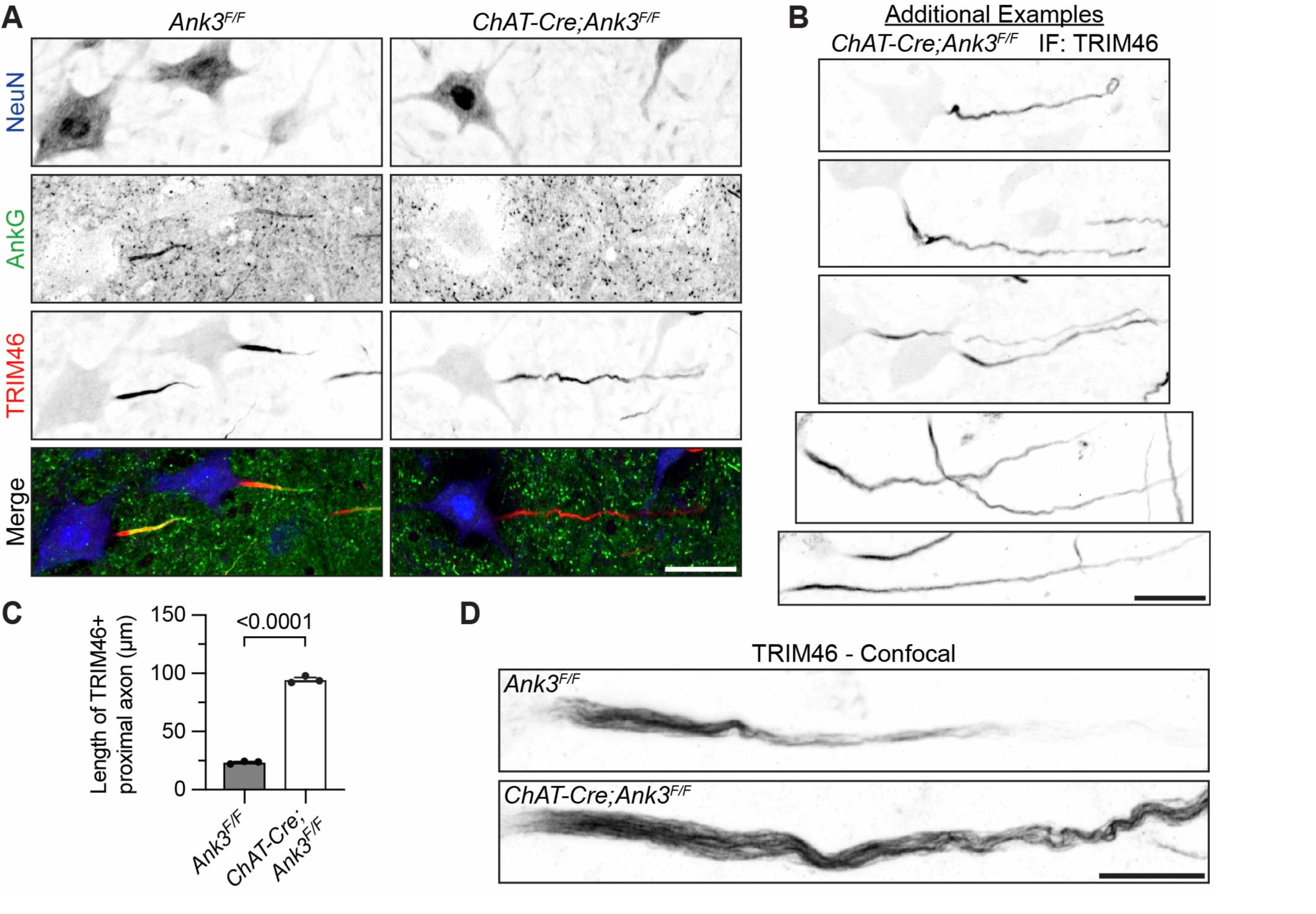
TRIM46 localization depends on AnkG *in vivo*. (A) Motor neurons in *Ank3^F/F^* and *ChAT-Cre;Ank3^F/F^* spinal cord immunostained for NeuN (blue), AnkG (green), and TRIM46 (red). (B) Examples of TRIM46 immunostaining in *ChAT-Cre;Ank3^F/F^* spinal cord. (C) Quantification of the length of proximal axon immunopositive for TRIM46 (*Ank3^F/F^* mean=23.52 μm, *ChAT-Cre;Ank3^F/F^* mean= 94.58 μm, N=26-52 axons per mouse, n=3 mice). Analyzed with unpaired t-test. (D) Confocal microscopy of TRIM46 in *Ank3^F/F^* and *ChAT-Cre;Ank3^F/F^* spinal cord. Error bars represent SEM. Scale bars: (A, B) 30 μm, (C) 10 μm.

To determine whether AnkG loss impacts TRIM46’s microtubule association in the proximal axon, we performed confocal imaging of the same tissues. TRIM46 exhibited a fibrillar staining pattern suggestive of microtubule binding in *ChAT-Cre;Ank3^F/F^* neurons as in control neurons (Fig. 8D). Whether the AnkG-deficient neurons still exhibit microtubule fascicles with cross-bridges, and, if so, whether fasciculation extends abnormally far down the axon alongside TRIM46, is unknown. A prior *in vivo* study found that AnkG-deficient Purkinje cells lack thick microtubule bundles, however, this study examined AIS by transmission electron microscopy only in the longitudinal plane and did not determine whether microtubule cross-bridging, which is rarely visible in this plane, was affected (Sobotzik et al., 2009). Thus, the precise state of microtubule fasciculation in AnkG-deficient axons remains unclear.

### AnkG and TRIM46 are independently detergent insoluble

The AIS is highly detergent-insoluble (Huang et al., 2017). This is thought to be due to AIS-resident proteins’ strong association with AnkG and other AIS cytoskeletal proteins (e.g. spectrins). To determine if AnkG remains detergent-insoluble in the absence of TRIM46, we used CRISPR to knock out TRIM46 in cultured rat hippocampal and cortical neurons. We used AAV to deliver control or TRIM46-specific CRISPR guide RNA (gRNA) and Cas9 constructs.

Neurons transduced with the TRIM46-specific gRNAs produced TRIM46-negative neurons (Fig. 9A, arrows). The percentage of neurons with TRIM46-positive AIS was roughly 90% lower in cultures treated with TRIM46 gRNA than in those treated with control gRNA, demonstrating the efficiency of the knockout (Fig. 9A-B). TRIM46 knockout did not significantly change the percentage of neurons with AnkG-positive AIS (Fig. 9C). After 30 minutes of solubilization with 0.5% Triton X-100 before fixation, MAP2 immunoreactivity was significantly reduced in all neurons. TRIM46 AIS staining in non-transduced neurons remained strong (Fig. 9D, arrowheads). Immunostaining revealed strong AnkG signal at the AIS of both TRIM46-positive (Fig. 9D, arrowheads) and TRIM46-negative neurons (Fig. 9D, arrows) after 30 minutes of solubilization. Next, to determine if the mislocalized TRIM46 observed in AnkG-deficient neurons (Fig. 8) remains detergent insoluble, we employed a similar CRISPR approach to knock out AnkG in cultured rat hippocampal neurons. TRIM46 proved detergent-insoluble, both when properly localized to the AIS of AnkG-positive neurons (Fig. 9E, arrowheads) and when mislocalized in the proximal axon of AnkG-negative neurons (Fig. 9E, arrows). Thus, AnkG and TRIM46 are both detergent-insoluble at the AIS/proximal axon and neither protein’s insolubility depends on the other’s.

**Fig. 9.**
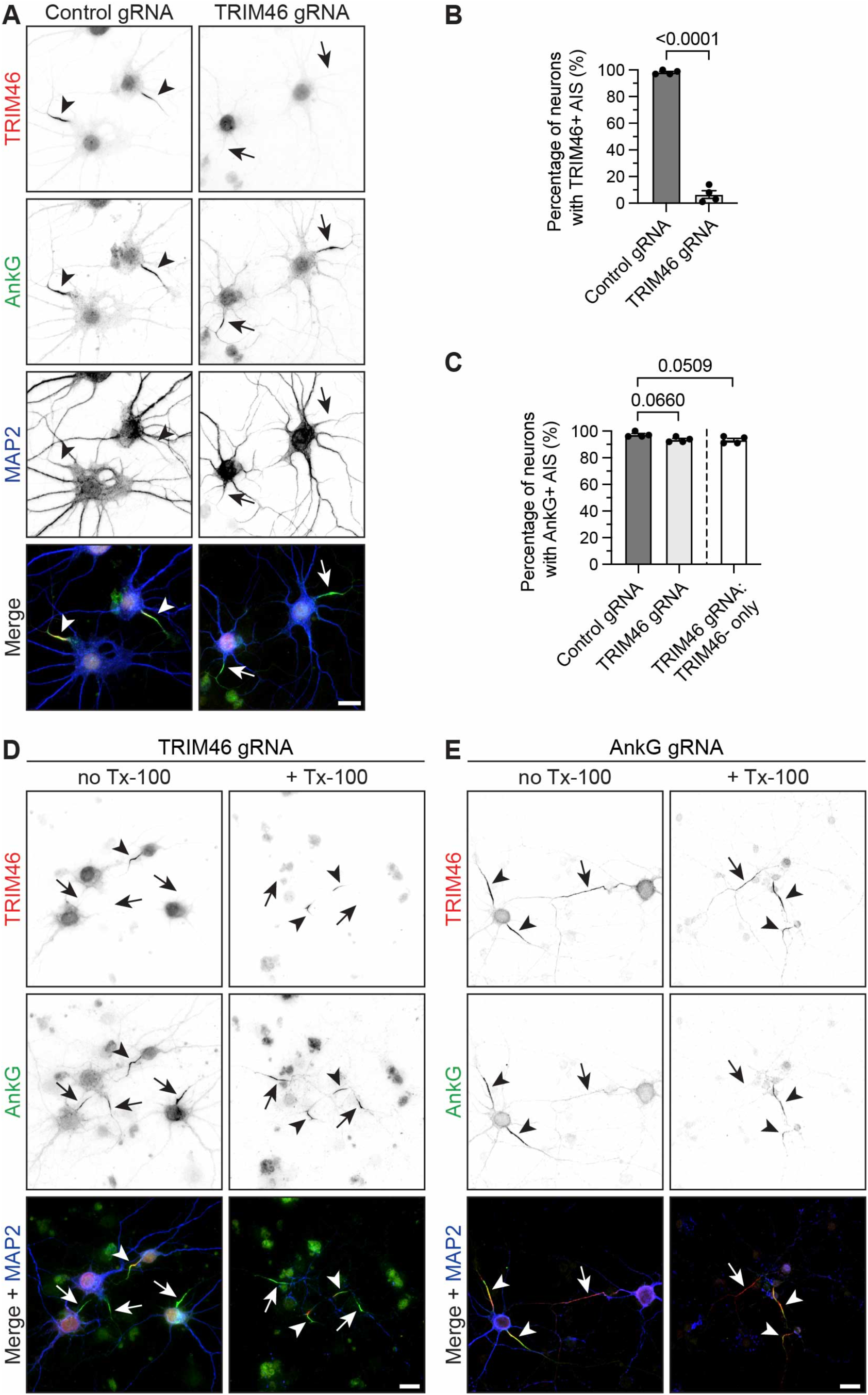
AnkG and TRIM46 are independently detergent insoluble. (A) Immunostaining for TRIM46 (red), AnkG (green), and MAP2 (blue) in cultured rat hippocampal neurons transduced with Cas9 and control or TRIM46 gRNA at DIV 0 and fixed at DIV 21. (B-C) Quantification from cultured rat cortical and hippocampal neurons transduced with Cas9 and control or TRIM46 gRNA between DIV 0-DIV 5 and fixed at DIV 17-DIV 21 (n=4, N=195-266 neurons pooled from 2-4 coverslips per replicate). (B) Percentage of neurons with a TRIM46-positive AIS in control (mean: 98.25%) and TRIM46 (mean: 6.5%) gRNA neurons. Analyzed with unpaired t-test. (C) Percentage of neurons with an AnkG-positive AIS in control (mean: 97.25%) and TRIM46 (mean: 93.5%) gRNA cultures. On the right, only TRIM46-negative neurons in the TRIM46 gRNA cultures are included (mean: 93.25%; N=193-250 neurons per replicate). Analyzed with ordinary one-way ANOVA (P=0.0501) with Dunnett’s multiple comparisons test (adjusted p-values shown on graph). (D) Immunostaining for TRIM46 (red), AnkG (green), and MAP2 (blue, shown only in the merged images) in cultured rat cortical neurons that were transduced with Cas9 and TRIM46 gRNA at DIV 1 and fixed at DIV 21. The neurons on the right were treated with Triton X-100 before fixation. Each field of view includes both TRIM46-negative (arrows) and TRIM46-positive (arrowheads) neurons for comparison purposes. (E) Immunostaining for TRIM46 (red), AnkG (green), and MAP2 (blue, shown only in the merged images) in cultured rat hippocampal neurons that were transduced with Cas9 and AnkG gRNA at DIV 0 and fixed at DIV 21. The neurons on the right were treated with Triton X-100 before fixation. Each field of view includes both AnkG-negative (arrows) and AnkG-positive (arrowheads) neurons for comparison purposes.

### TRIM46 is found at proximal nodes of Ranvier

Although microtubule fasciculation is usually discussed in relation to the AIS, it has also been observed at some nodes of Ranvier that are near to the cell bodies of motor and sensory neurons of the spinal cord and dorsal root ganglia (DRG) (Nakazawa and Ishikawa, 1995).

TRIM46 was later observed at nodes within the DRG (Gumy et al., 2017; Nascimento et al., 2022). To determine if most nodes have TRIM46, we immunolabeled TRIM46 in preparations from wildtype mice comprised of the sciatic nerve with several attached DRGs and dorsal and ventral roots. In this preparation, proximal nodes are found in three places: proximal nodes of motor neurons are located at end of the ventral root farthest from the DRG (nearest the spinal cord *in situ*), while proximal nodes of sensory neurons are found on both sides of the DRG, with the central nodes being in the dorsal root projecting to the spinal cord and the peripheral nodes on the side of the sciatic nerve. We observed nodal TRIM46 in each of these locations, but not in any of their more distal counterparts (Fig. 10A). TRIM46 extends beyond the AnkG-labeled nodes and into the flanking Caspr-labeled paranodes, showing that, as at the AIS, it is not strictly colocalized with AnkG (Fig. 10B, arrowheads). These findings confirm that TRIM46 is found in proximal nodes of spinal cord motor neurons and both the CNS and PNS branches of DRG sensory neurons, which are the only known sites of neuronal microtubule fasciculation besides the AIS.

**Fig. 10.**
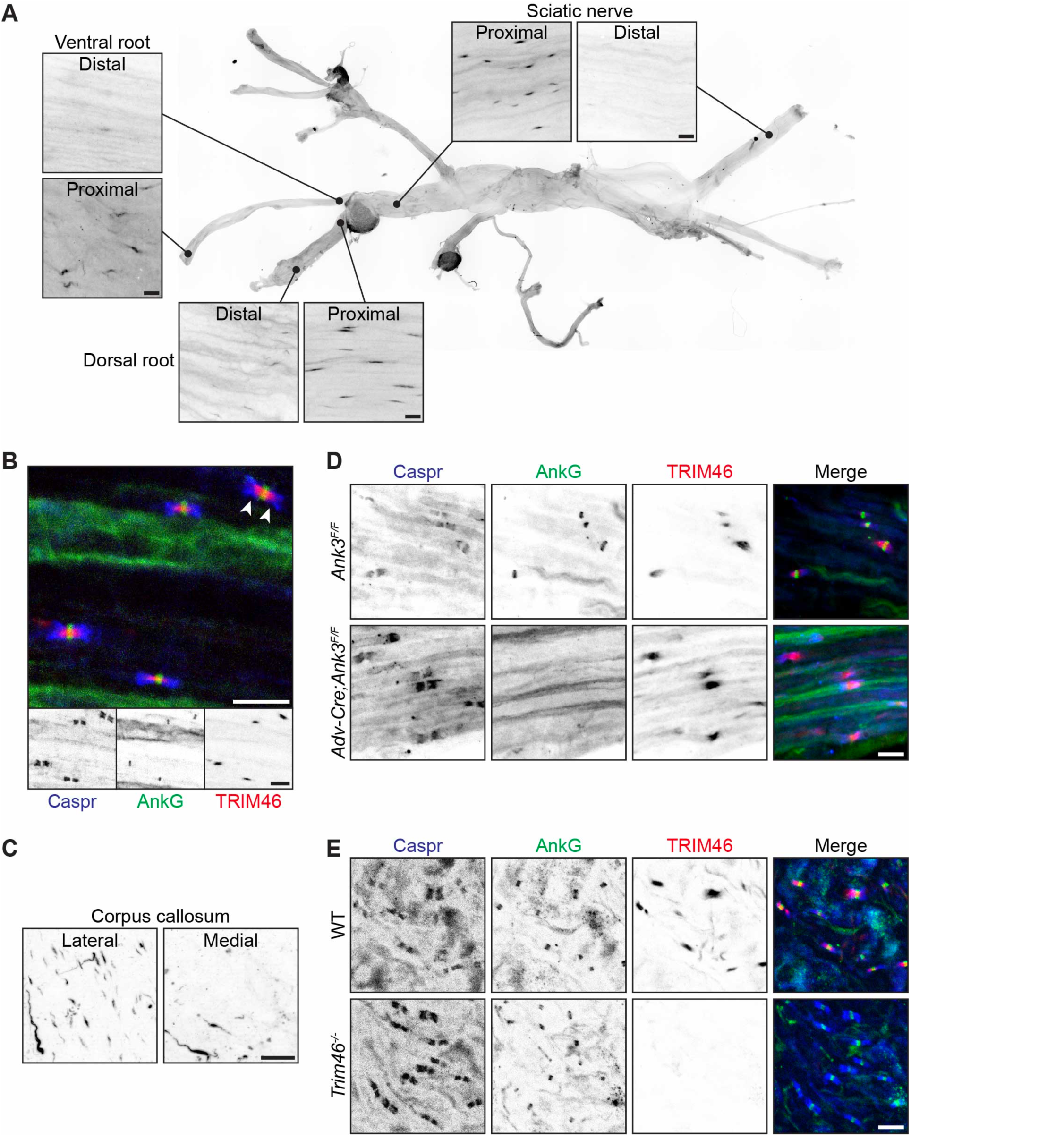
TRIM46 is found at proximal nodes of Ranvier. (A) TRIM46 immunostaining in wildtype adult sciatic nerve, ventral root, and dorsal root. (B) Immunostaining for Caspr (blue), AnkG (green), and TRIM46 (red) in wildtype proximal sciatic nerve. (C) TRIM46 immunostaining in wildtype corpus callosum. (D-E) Immunostaining for Caspr (blue), AnkG (green), and TRIM46 (red) in (D) dorsal root from adult *Ank3^F/F^* and *Adv-Cre;Ank3^F/F^* mice and (E) DRG from adult wildtype and *Trim46^-/-^* mice. Scale bars: 10 μm.

To determine if nodal TRIM46 is found in the brain, we immunostained wildtype corpus callosum for TRIM46 (Fig. 10C). We reasoned that in coronal brain sections the most lateral, ventral edge of the corpus callosum would contain a higher proportion of nodes that are close to cortical neuron cell bodies than would the medial, dorsal region that bridges the hemispheres. In comparing these two regions, we observed nodal TRIM46 in the lateral but not medial corpus callosum. These results show that TRIM46 is found at proximal nodes throughout the nervous system, suggesting that microtubule fasciculation is a common feature of all proximal nodes.

After finding TRIM46 at nodes, we examined the relationship between nodal TRIM46 and AnkG. To determine whether loss of AnkG affects nodal TRIM46, we immunostained DRGs and dorsal roots from *Adv-Cre;Ank3^F/F^* mice. These mice lack AnkG only in Advillin-expressing cells such as DRG neurons. We observed that TRIM46 is still present at nodes lacking AnkG (Fig. 10D). In these nodes, we did not see a mislocalization of TRIM46 like that found in AnkG-null proximal axons (Fig. 8). In AnkG-null neurons, ankyrinR can compensate and assemble nodes but not the AIS (Ho et al., 2014; Liu et al., 2020), which may explain the redistribution of TRIM46 at proximal axons but not nodes lacking AnkG. To determine whether loss of TRIM46 affects nodal AnkG, we immunostained *Trim46^-/-^* DRGs and observed that AnkG is still clustered at nodes lacking TRIM46 (Fig. 10E). We conclude that TRIM46 is not required for the formation of AIS or nodes *in vivo*.

## DISCUSSION

The first loss-of-function study of neuronal TRIM46 used shRNA to silence TRIM46 expression in cultured neurons. That study reported that the loss of TRIM46 reduced AnkG-labeled AIS by ∼60%, leading to the conclusion that TRIM46 is necessary for AIS assembly (van Beuningen et al., 2015). This claim was partially supported by a second *in vitro* study, where neurons derived from TRIM46-null mouse embryonic stem cells showed a more modest 35% reduction in the percentage of neurons with an AnkG-positive AIS (Vuong et al., 2022). Recently, Guan et al. (2024) reported a TRIM46 knockout rat model and measured a ∼30% reduction in the percentage of brain cells with an AIS. In contrast to these previous studies, our results obtained from rigorously validated *Trim46^-/-^* mice showed neither a reduced percentage of neurons with AIS proteins nor a reduction in the fluorescence intensity of AnkG. We conclude that TRIM46 is not required for axon specification or AIS function, assembly, or maintenance *in vivo*.

Why have different studies of TRIM46 loss produced opposing conclusions about the necessity of TRIM46 for AIS formation? The difference may primarily reflect the models used to study TRIM46 function. For example, it is possible the *Trim46^-/-^* mice we used are not a bona fide knockout. However, we showed that *Trim46^-/-^* mice express neither wildtype TRIM46 nor a mutant N- or C-terminal fragment. It is possible that the *Trim46^-/-^* mice express a variant that skips the epitopes of all the antibodies we used, either a previously unknown isoform or a *de novo* mutant. However, we consider these possibilities unlikely, since the four antibodies we used have epitopes spanning three of TRIM46’s ten exons, one of which also contains the RING finger domain previously shown to be required for TRIM46’s AIS localization (van Beuningen et al., 2015). Considering this and our validation data, the simplest conclusion is that *Trim46^-/-^* mice lack TRIM46 protein.

Alternatively, the effects of TRIM46 knockdown might be explained by toxic or off-target effects of the shRNA rather than depletion of TRIM46 itself. However, transfection with TRIM46 rescues the effects of TRIM46 shRNA (van Beuningen et al., 2015; Harterink et al., 2019), suggesting those effects were indeed caused by TRIM46 loss. That AIS formation was also diminished in cultured TRIM46 knockout neurons further suggests a role for TRIM46 in AIS formation (Vuong et al., 2022). It is also possible that compared to the intact brain environment, cultured neurons may be more susceptible to the loss of TRIM46. We previously showed that AIS are highly susceptible to disruption by injury, or any condition that results in elevated intracellular calcium and subsequent proteolysis of AIS proteins (Schafer et al., 2009).

Despite two independent *in vitro* lines of evidence for TRIM46’s involvement in AIS formation, our *in vivo* study demonstrates that it is dispensable in the brain. A compensatory mechanism analogous to AnkR’s role at nodes of Ranvier could reconcile these apparently conflicting findings. Nodes usually lack AnkR and are assembled by AnkG; however, when AnkG is absent, AnkR can compensate (Ho et al., 2014; Liu et al., 2020). Thus, considering node assembly strictly in terms of necessity paints an incomplete picture—to describe how wildtype nodes form, a major role must be credited to AnkG even though it is not actually required. In the same way, it may be that TRIM46 is involved in AIS formation when it is present, but when absent another protein or mechanism can compensate. Future studies will test this hypothesis by searching for compensatory mechanisms in *Trim46^-/-^* mice.

If compensation explains the preserved AIS formation in *Trim46^-/-^* mice, why were AIS not rescued in the prior *in vitro* studies? This may reflect differences in the timing of experimental manipulation and analysis. In the shRNA experiments that produced the first and most extreme indications of TRIM46’s AIS function, wildtype neurons underwent some degree of normal axon development and specification before being abruptly depleted of TRIM46 and then evaluated within a few days of that loss (van Beuningen et al., 2015). As noted by Vuong et al. (2022), the sudden loss of TRIM46 after it had already begun to function at the AIS might have more severe consequences than would be seen in neurons that never express any TRIM46 where compensatory mechanisms may be engaged earlier. Indeed, cultured neurons derived from TRIM46 knockout ESCs (Vuong et al., 2022) showed a milder impairment of AIS formation than TRIM46 knockdown neurons (van Beuningen et al., 2015). In addition, the knockout neurons were evaluated at DIV 7 while the knockdown neurons were evaluated at DIV 4, a relatively early time when not all neurons have an AIS (Yang et al., 2007). Thus, the knockout neurons *in vitro* both lost TRIM46 earlier and were evaluated at a more mature stage than the knockdown neurons. These experimental differences may explain the varying results. Together, these results suggest that AIS formation is less impaired in neurons that have more opportunity to adjust to TRIM46 loss.

Our results in *Trim46^-/-^* mice, which never express TRIM46 protein and can be studied over a much longer period than cultured neurons, show no reduction in the number of AIS, length of the AIS, or AnkG fluorescence intensity; axons are also appropriately specified. This is in contrast to the results of Guan et al. (2024) who reported a ∼30% reduction in the percentage of brain cells with AIS in a TRIM46 knockout rat model. Again, there are notable differences in the experimental approaches they used to study AIS compared to our methods. For example, Guan et al. (2024) did not fix the brain until after sectioning. This differs from the more common methods of AIS immunostaining where tissues are lightly fixed prior to sectioning. The AIS is extremely sensitive to both overfixation and proteolysis by endogenous calpains (Schafer et al., 2009); the lack of rapid fixation to quench calpain activity and postfixation of thin brain slices can reduce antigenicity. Consistent with these possibilities, the immunostaining for AnkG produced fractured and irregular AIS staining in addition to the reported loss of AIS staining.

While we show that TRIM46 is dispensable for AIS formation *in vivo*, we did not determine whether *Trim46^-/-^* AIS have microtubule fascicles due to the technical challenges of observing cross-linked microtubules at the AIS by electron microscopy. Prior studies reported that TRIM46 is localized at AIS microtubule fascicles in cultured neurons, that TRIM46 is necessary for their formation, and that TRIM46 is sufficient to induce microtubule fasciculation in HeLa cells (van Beuningen et al., 2015; Harterink et al., 2019). It will be important in future studies to evaluate whether microtubules still form fascicles in *Trim46^-/-^* mice. It is possible that compensatory mechanisms that preserve AIS formation also preserve microtubule fasciculation, or that these phenomena are independent. With the observation that TRIM46 is present at proximal nodes of Ranvier in motor neurons, TRIM46 is now recognized at all known sites of neuronal microtubule fasciculation (Nakazawa and Ishikawa, 1995). We also found TRIM46 at proximal nodes in brain, suggesting that microtubule fasciculation in proximal nodes is ubiquitous. That TRIM46 is present at nodes of Ranvier and proximal axons in AnkG knockout neurons suggests that there are AnkG-independent mechanisms allowing TRIM46 to be localized at these sites.

Neuronal TRIM46 may have roles beyond AIS assembly that we did not address. For example, TRIM46 knockdown was reported to induce a mixed microtubule orientation in axons (where microtubules are normally uniformly plus-end-out), impair axonal transport, and disrupt the compartmentalization of somatodendritic and axonal proteins (van Beuningen et al., 2015; Fréal et al., 2019). Similar effects have been observed in AnkG-deficient neurons (Hedstrom et al., 2008; Fréal et al., 2016, 2019; Kuijpers et al., 2016; Teliska et al., 2022), so they could be secondary to the AIS disruption caused by TRIM46 knockdown, rather than direct results of TRIM46 loss. It will be interesting to evaluate these aspects of neuronal polarity in *Trim46^-/-^* mice, especially considering previous findings on neuronal polarity in AnkG-deficient mice and neurons (Hedstrom et al., 2008; Sobotzik et al., 2009).

In summary, this first study of *Trim46^-/-^* mice indicates that TRIM46 is not required for axon specification or AIS formation *in vivo*, possibly due to an unknown compensatory mechanism. Future study of this compensation will improve our understanding of how the AIS works with the microtubule cytoskeleton to support neuronal polarity and function.

## AUTHOR CONTRIBUTIONS

A. J. M., V. L. P., Y. O., and M. N. R. designed research, analyzed data, and wrote the paper. A. J. M., V. L. P, and Y. O. performed research.

## Acknowledgements

Supported by grants from the National institutes of Health: R35 NS122073 (M. N. R.) and F31 NS134125 (A. J. M.) and by the Dr. Miriam and Sheldon G. Adelson Medical Research Foundation (M. N. R.). We thank Dr. Lindsay Teliska for help with AnkG cKO mice and Dr. Xiaoyun Ding for help with behavioral experiments.

